# Anti-NS1 antibodies induced by immunization with dendritic cell targeted hybrid monoclonal antibodies do not induce platelet dysfunction

**DOI:** 10.1101/2020.09.10.292334

**Authors:** Elaine Cristina Matos Vicentin, Kelly Nazaré da Silva Amorim, Marcio Massao Yamamoto, Higo Fernando dos Santos Souza, Daniela Santoro Rosa, Antonio José da Silva Gonçalves, Ada Maria de Barcelos Alves, Silvia Beatriz Boscardin

## Abstract

Dengue fever causes a disease whose symptoms range from asymptomatic to more severe, that may even lead to death. During infection, the non-structural (NS) 1 protein is produced in infected cells and subsequently shed to the circulation. Anti-NS1 antibodies have been implicated in disease pathogenesis as well as in protection against lethal infection. Here we analysed the role of anti-NS1 antibodies elicited when the NS1 protein was specifically targeted to two distinct dendritic cell (DC) subsets by chimeric monoclonal antibodies (mAbs) that bind to the DC surface receptors DEC205 and DCIR2. BALB/c mice received two doses of αDEC-NS1 or αDCIR2-NS1 mAbs in the presence of an adjuvant. We observed that anti-NS1 antibodies induced by immunization with chimeric αDEC-NS1 or αDCIR2-NS1 mAbs did not cause significant pathology or were able to protect BALB/c mice from an intracranial DENV2 NGC strain challenge.

## Introduction

Dengue fever (DF) is the most common clinical manifestation of dengue virus (DENV) infection that can additionally lead to the more severe dengue haemorrhagic fever/dengue shock syndromes (DHF/DSS), characterized by vascular leakage and shock (24). There are four viral serotypes (DENV-1, −2, −3 and −4), and it is estimated that DENV infects 390 million people annually causing approximately 96 million symptomatic infections, and 20,000 deaths due to DHF/DSS (5). The pathophysiology of the disease is complex and still not completely understood, as infected individuals can be asymptomatic, or present classic DF symptoms that may or may not evolve to the potentially fatal DHF/DSS. It is well documented the correlation between the occurrence of a secondary heterologous infection with DENV and severe dengue (16). This phenomenon may be explained by the generation of serotype cross reactive T cells that induce an exacerbated, but ineffective, response during the secondary infection, and by antibody-dependent enhancement (ADE) where cross-reactive, non-neutralizing, antibodies against structural proteins enhance DENV infection of Fcγ bearing cells, leading to an increase in viral load (16, 20, 22).

The 10.7 kb DENV genome encodes three structural (capsid, membrane and envelope) and seven non structural (NS) proteins that are involved with virus replication and assembly. NS1 is a ∼48 to −55-kDa glycosylated protein that is produced early during infection, and detected in the blood of infected individuals as early as the first day of symptoms onset (18). The presence of NS1 in the circulation induces the production of high anti-NS1 antibody titres in the infected individuals, and the levels of NS1 circulating in the blood have been correlated with an increase in the disease severity. For that reason, different studies have focused on studying the capacity of anti-NS1 antibodies to bind to different cells. In fact, anti-NS1 antibodies were already shown to bind to platelets, endothelial cells, etc (8, 9, 14). It seems that the cross-reactive epitopes are present on the C-terminal portion of NS1. When this region was removed or substituted by the C-terminal portion of the Japanese encephalitis virus, binding was abrogated (7). Anti-NS1 antibodies seem to bind to the WCxxSCxLxP motif on the protein disulfide isomerase on the surface of human platelets (8), increase bleeding time, and inhibit ADP mediated platelet activation and aggregation (7). Based on this evidence, anti-NS1 antibodies, especially those directed against the C-terminal portion of the molecule, have been implicated in the pathogenesis of the disease. On the other hand, NS1, but not anti-NS1 antibodies, has been directly implicated in dengue pathogenesis. Two recent papers elegantly showed that NS1 is able to activate TLR4, induce the production of pro-inflammatory cytokines and cause endothelial cell leakage that is a hallmark of DHF/DSS (21). Also, the administration of the hexameric form of NS1 together with a sub lethal dose of DENV2 increased the death rate of mice and produce high vascular leakage. Such leakage was reverted in the presence of anti-NS1 antibodies (4). More studies are required to examine the real contribution of NS1 or anti-NS1 antibodies to the development of severe dengue symptoms.

In contrast to its role in the development of disease pathogenesis, NS1 has also been used in immunization protocols as a vaccine candidate. Partial protection against disease has been shown in a mouse model of intracranial virus administration (1, 10, 17).

Our group has been developing an immunization strategy that targets NS1 directly to dendritic cells by using chimeric monoclonal antibodies (mAbs) directed against DC endocytic surface receptors fused with NS1 (17). We used two mAbs that specifically directed the NS1 protein to two different DCs subsets known as CD8α+ and CD8α-DCs (13). CD8α+ DCs subset expresses the DEC205 endocytic receptor while CD8α-DCs express another receptor known as DCIR2. NS1 targeting to the CD8α+DEC205+ DC subset was able to induce T cells that were responsible for the partial protection observed after challenge of immunized mice with the DENV-2 NGC strain. Interestingly, anti-NS1 antibody titres were very similar when NS1 was targeted to the CD8α+DEC205+ or the CD8α-DCIR2+ DC subsets, and did not seem to play any role in protection after challenge (17). These results prompted us to explore in a little more detail the effects of anti-NS1 antibodies in disease pathogenesis. Based on the information provided above, we set out to investigate if the high anti-NS1 antibody titres could be detrimental to platelet function. Here we analyse the effect of anti-NS1 antibodies elicited by immunization of BALB/c mice with αDEC-NS1 or αDCIR2-NS1 hybrid mAbs in platelet function, and also in protection during a lethal challenge.

## Materials and Methods

### Mice

Six-to 8-week-old male BALB/c mice were bred at the Isogenic Mouse Facility of the Parasitology Department, University of São Paulo, Brazil. All animal experiments complied with the ARRIVE guidelines and with the recommendations of the National Institutes of Health guide for the care and use of laboratory animals and the Brazilian National Law (11.794/2008). The Institutional Animal Care and Use Committee (CEUA) of the University of São Paulo (protocol number 082) and the Animal Use Ethical Committee of Fundação Oswaldo Cruz (approval ID: L-067/08) approved all protocols.

### Chimeric monoclonal antibody production

The chimeric αDEC-NS1 and αDCIR2-NS1 mAbs, together with empty αDEC and αDCIR2, were produced exactly as described in (17). After purification, chimeric mAbs were dialyzed against cold PBS, filtered through 0.2 mM membranes (TPP), and ran in 12% SDS-PAGE gels under denaturing conditions to check for degradation. Chimeric mAbs were maintained at −20 °C until use.

### Immunizations and immune sera obtainment

Groups of 5-10 mice were immunized intraperitoneally (i.p.) with 2 doses (30-day interval) of 5 μg of each mAb (αDEC-NS1, αDCIR2-NS1, αDEC or αDCIR2) together with 50 μg poly (I:C) (Invivogen), exactly as described in (17). A control group was also immunized with 10 μg of recombinant NS1 protein produced as previously described (2) plus poly (I:C). Fourteen days after the administration of the second dose, mice were bled and serum was obtained. Sera were then pooled for each group and IgGs were purified using protein G beads (GE Healthcare). The IgG concentration for each sample was obtained measuring the absorbance at 280 nm using a nanodrop (Thermo Scientific).

### Analysis of NS1-specific antibodies

For the detection of NS1-specific antibodies, ELISA assays were performed exactly as described previously (6). The avidity index was calculated using an extra step of incubation with 7M urea for 5 min, as described in (12, 25).

### Platelet preparation and binding assay

Platelets were obtained from 3 healthy volunteers that provided written informed consent. The protocol was approved by the Ethics Committee of the Institute of Biomedical Sciences, University of São Paulo, Brazil (protocol number 26301814.4.0000.5467).

Platelet preparation and antibody binding were performed for each individual volunteer as described in (7). Briefly, 20 mL of whole human blood containing 2 mL of anticoagulant (29.9 mM sodium citrate, 113.8 mM glucose, 72.6 mM NaCl, and 2.9 mM citric acid, pH 6.4) were centrifuged at 100 xg for 20 min at room temperature. Platelet rich plasma was then removed and centrifuged at 1,000 xg for 10 min at room temperature. Platelets were then washed twice with PBS containing 50 mM EDTA and suspended in Tyrode’s solution (137 mM NaCl, 20 mM HEPES, 3.3 mM NaH_2_PO_4_, 2.7 mM KCl, 1 mg/ml BSA, and 5.6 mM glucose, pH 7.4) at a concentration of 10^6^ platelets/ml. After fixation with 1% formaldehyde in PBS at room temperature for 10 min, platelets were washed with PBS and incubated for 30 min at room temperature with 2.5 or 5 μg of the purified IgGs from each immunized group. After two additional washes, samples were incubated with goat Alexa488-conjugated anti-mouse IgG (Jackson ImmunoResearch Laboratories) for 30 min and washed twice. Three hundred thousand events were collected inside the platelet gate using a FACS Canto flow cytometer (BD Biosciences). Analysis was performed using FlowJo software (version 9.3, Tree Star, San Carlo, CA).

### Bleeding time and prothrombin time

Bleeding tendency was assessed in each group of immunized animals (n= 7-10) on days 1, 3 and 7 after the second immunization. We used Whatman filter paper strips, in which the transected tail tip (3mm) was gently touched onto filter paper immediately, and after every 30 second interval for the first 3 min. If after that time the tail was still bleeding, the drops were collected every 10 seconds until bleeding stopped.

Prothrombin time test was performed using the Soluplastin kit (Wiener Lab, Argentina), according to the manufacturer’s instructions. Mice were bled individually 7 days after the administration of the second dose for plasma obtainment.

### Platelet counts

Whole blood samples were obtained from individual mice 7 days after the second dose and the platelet numbers were analysed using the BC-5300 VET Auto hematology analyser (Mindray, China).

### ADP-stimulated platelet aggregation assay

Human whole blood was mixed with 0.11 M of the anticoagulant sodium citrate at a proportion of 9:1 and centrifuged at 100 xg for 10 min at room temperature. The platelet-rich plasma (PRP) was collected from the upper layer while the lower layer was subsequently centrifuged at 1000 xg for 10 min and the supernatant was collected as platelet-poor plasma (PPP). The PPP was used as blank for analysis of platelet aggregation. The PRP was incubated with 25 μg of the purified IgGs derived from the different groups or control mouse at 37 °C for 30 min followed by addition of ADP (10μM final concentration). Platelet aggregation was detected using an automated aggregometer (Chrono-Log Corporation, Model 700).

### Purified IgG transfer and challenge

Groups of 10 naïve BALB/c mice were transferred i.p. with 100 μg of IgGs purified as described above and administered twice, one day before and three days after challenge. For the challenge assays, animals were anesthetized with a mixture of ketamine-xylazine and then inoculated by the intracranial route (i.c.) with 30 µL of DENV2 NGC strain corresponding to approximately 40 LD50 (11). After challenge, mice were monitored daily for 21 days for signs of morbidity according to the scale described in (3). After that period, animals that survived challenge were euthanized.

### Statistical analysis

Statistical significance was calculated using one-way ANOVA followed by Tukey’s honestly significantly different (HSD). Log-rank test was used to compare mouse survival times after challenge. All differences were calculated using Prism 5 software (GraphPad Software Inc, La Jolla, CA). Differences with p<0.05 were considered statistically significant.

## Results

### Purified IgGs from αDEC-NS1, αDCIR2-NS1 or recombinant NS1 protein immunized mice recognize the recombinant protein

To study in more detail the anti-NS1 antibodies elicited by the immunization with αDEC-NS1, αDCIR2-NS1 hybrid mAbs or with the recombinant NS1 protein (rec. NS1) in the presence of poly (I:C), we purified total IgGs from the sera of BALB/c immunized mice. As negative controls, groups of animals were also immunized with αDEC or αDCIR2 mAbs that did not contain any fused protein. Figure 1A shows a reduced SDS-PAGE containing 1 μg of total IgGs purified from the different groups. In all purifications we identified 50 kDa heavy chains and 25 kDa light chains. As expected, IgGs obtained from the groups immunized with αDEC-NS1, αDCIR2-NS1 hybrid mAbs or with rec. NS1 recognized the NS1 protein adsorbed in ELISA plates. A binding curve was constructed using different concentrations of total IgGs (figure 1B). As we can observe, mice immunized with αDEC-NS1 or αDCIR2-NS1 hybrid mAbs showed a higher concentration of anti-NS1 antibodies when compared to the group immunized with the rec. NS1. We next sought to measure the avidity index of the anti-NS1 antibodies induced in mice immunized with the αDEC-NS1, αDCIR2-NS1 hybrid mAbs or with rec. NS1. The results indicated that the anti-NS1 antibody avidity index was lower in the sera of mice immunized with αDEC-NS1 when compared to anti-NS1 antibodies induced in mice immunized with either αDCIR2-NS1 mAbs or with the rec. NS1. Interestingly, the higher avidity index was observed in mice immunized with rec. NS1 (figure 1C).

**Figure 1.**
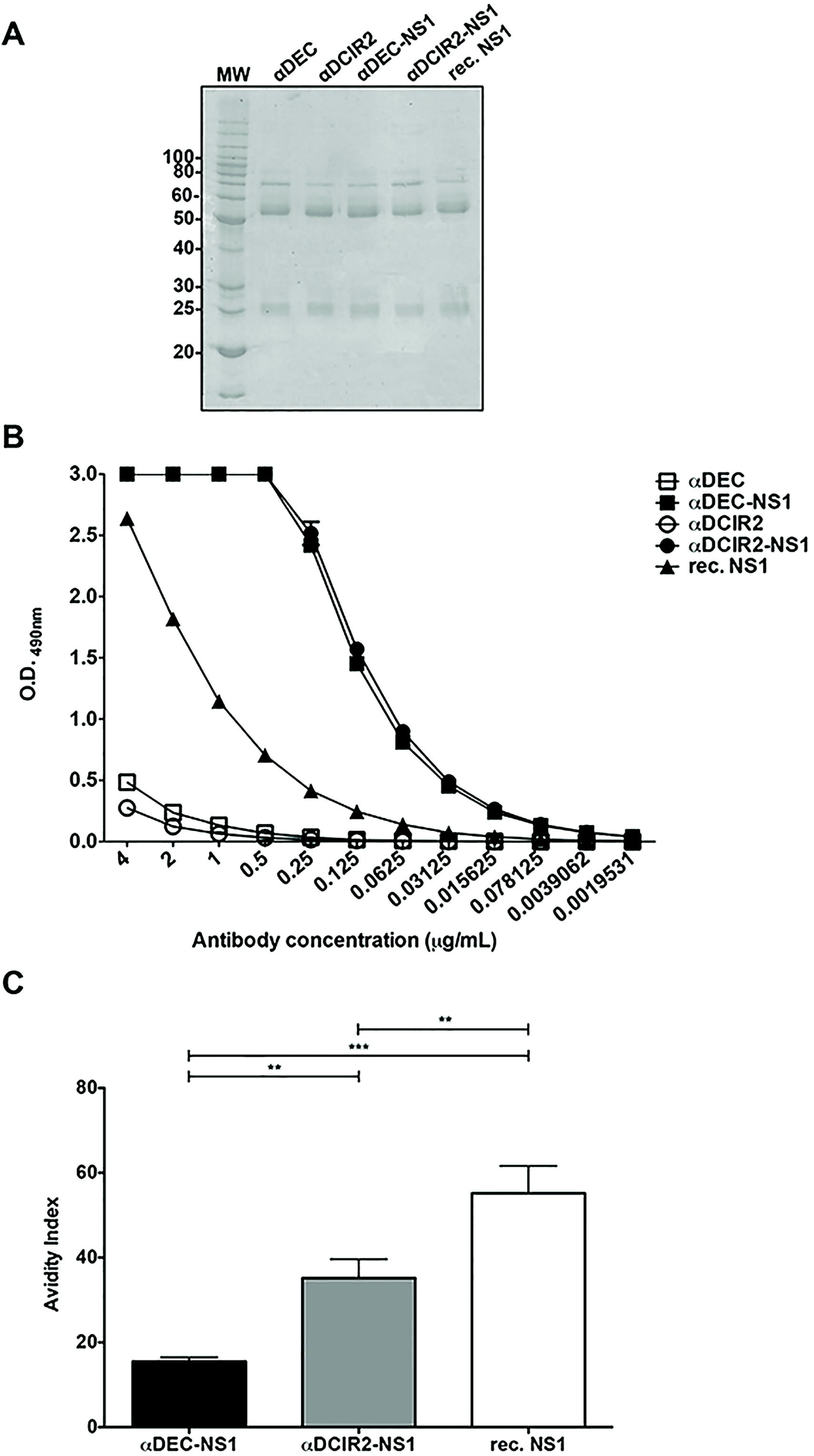

### Evaluation of the capacity of purified IgGs to bind to platelets and block their aggregation

Previous evidence indicated that anti-NS1 antibodies generated during dengue infection can cross react with epitopes present on human blood clotting pathway (14) and inhibit platelet aggregation (19). To verify if the anti-NS1 antibodies induced in mice immunized with the hybrid mAbs would cross-react with platelets, we incubated human platelets with different amounts of IgGs purified from the sera of the different groups. Figure 2A shows the binding curves observed when 5 µg of the purified antibodies from the different groups were incubated with the platelets. Figure 2B presents the percent of positive cells when 2,5 or 5 µg of the purified antibodies derived from each group were incubated with the platelets. The readout was obtained using an anti-mouse-IgG antibody. Finally, figure 2C shows the median fluorescence obtained using the same methodology. Serum samples from mice immunized with the αDCIR2-NS1 mAb reacted more strongly with human platelets when compared to sera harvested from mice immunized with the αDEC-NS1 mAb or from the controls (αDEC or αDCIR2, p<0.001). We also examined the effect of anti-NS1 Abs on ADP-induced platelet aggregation. Results showed no statistically significant difference among the different groups (p>0.05), indicating that the anti-NS1 antibodies elicited by the immunization with αDEC-NS1, αDCIR2-NS1 or rec. NS1 are not able to block ADP-induced platelet aggregation (figure 3). So, despite the binding of the anti-NS1 antibodies to platelets observed with the purified IgGs from animals immunized with αDCIR2-NS1 (figure 2), these antibodies were not able to block platelet aggregation (figure 3).

**Figure 2.**
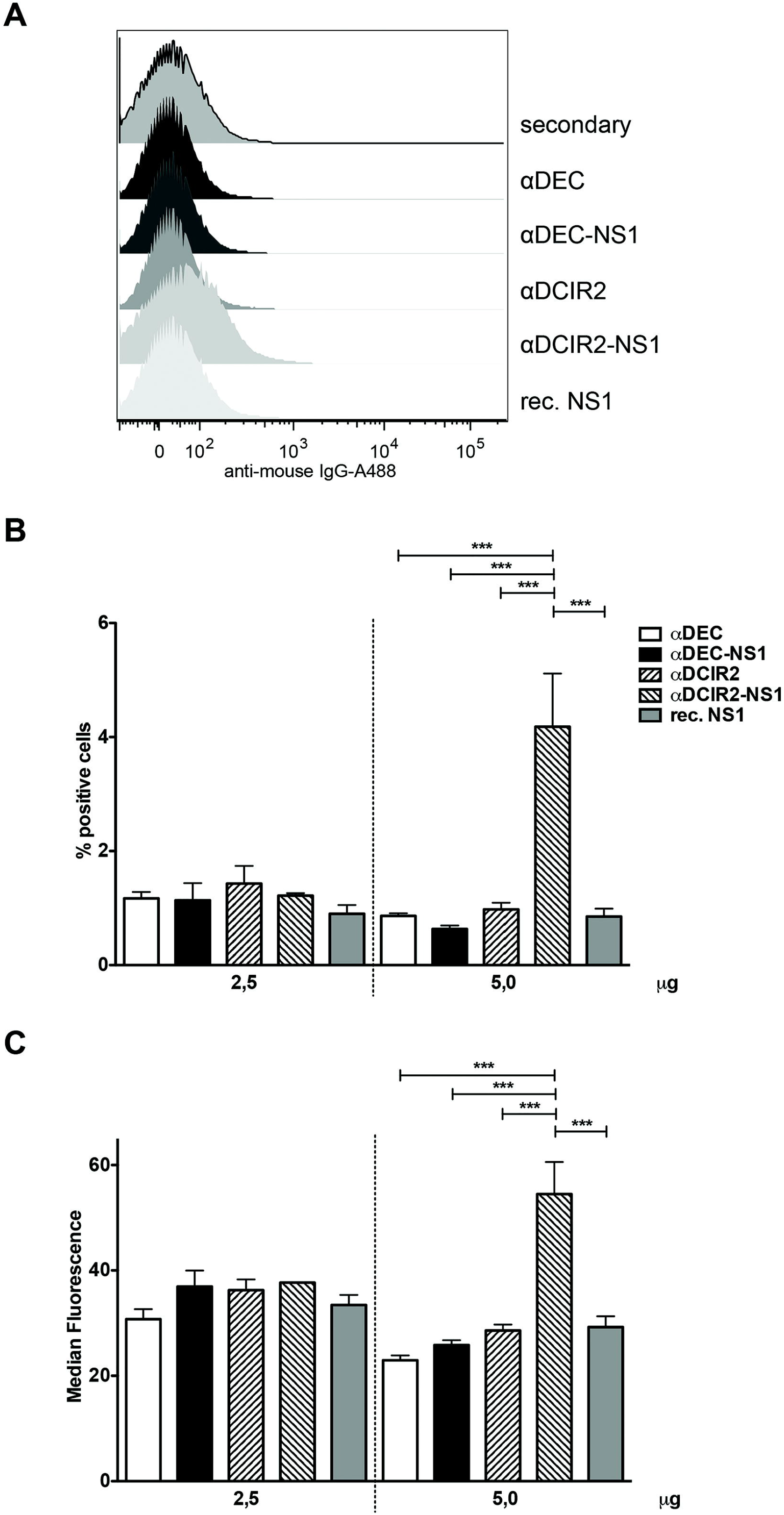

**Figure 3.**
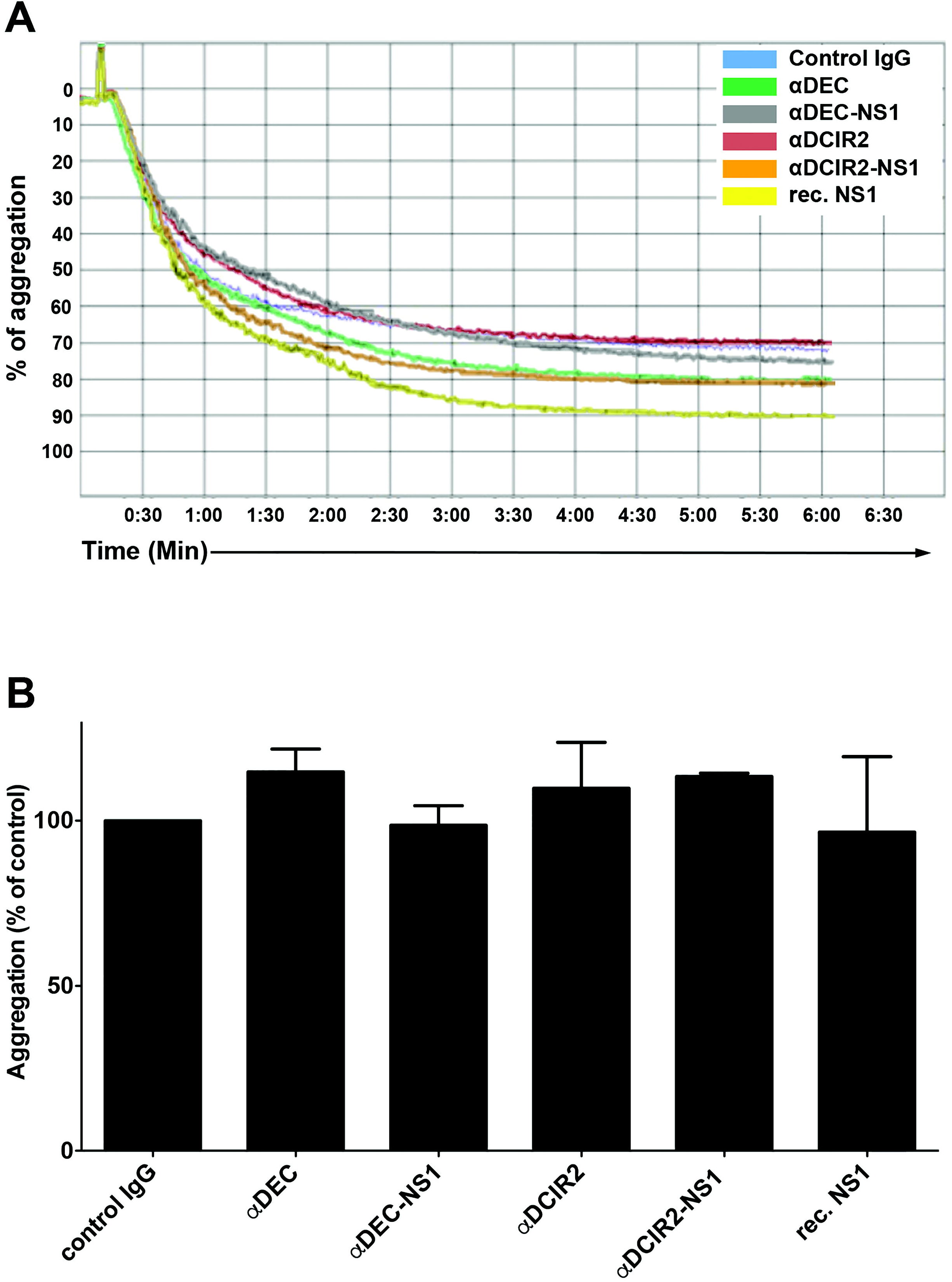

### Immunization with hybrid αDEC-NS1, αDCIR2-NS1 or rec. NS1 in the presence of poly (I:C) does not alter platelet numbers, prothrombin time or bleeding time

We next investigated if the immunization would alter bleeding time, platelet numbers or prothrombin time. For that purpose, groups of BALB/c mice were immunized twice (as described in Materials and Methods) with αDEC-NS1, αDCIR2-NS1 or rec. NS1 and with αDEC or αDCIR2, as controls. No statistically significant differences were observed in bleeding time on days 1, 3 or 7 after the second immunization (p>0.05; figure 4A). In addition, platelet numbers and prothrombin were also not different among the groups on day 7 after boost (p>0.05; figure 4B and 4C). In summary, these results indicate that our immunization regimen with either hybrid mAbs containing NS1 or with the rec. NS1 do not significantly alter the coagulation pathway.

**Figure 4.**
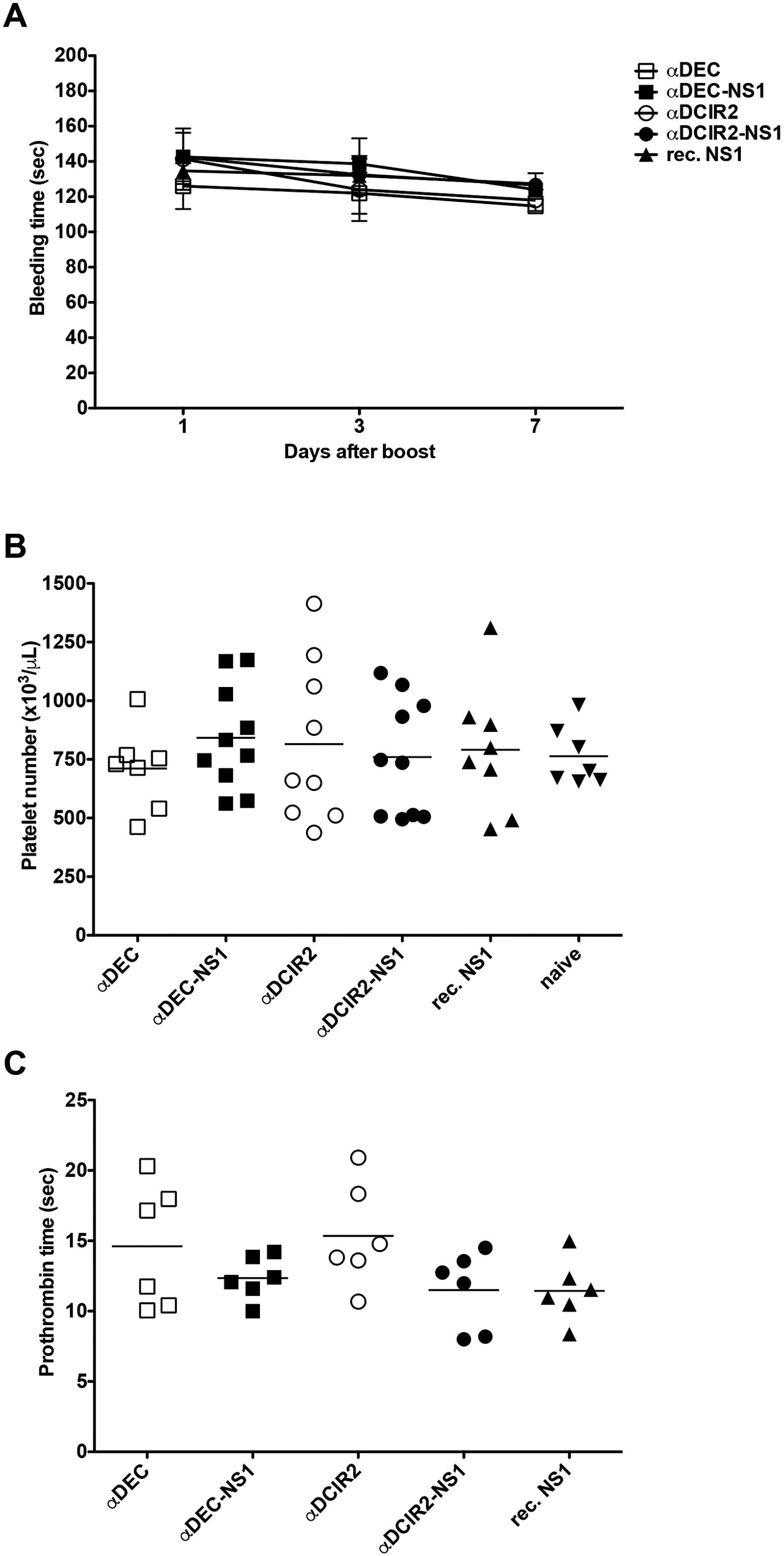

### Anti-NS1 rich IgGs do not protect mice against challenge with the DENV2 NGC strain

We have previously shown that CD4^+^ and CD8^+^ T cells elicited in mice immunized with the αDEC-NS1 mAb were partially protective against a lethal challenge with DENV2 NGC strain (17). However, the role of the anti-NS1 antibodies was not directly evaluated. For that purpose, we immunized mice with the different hybrid mAbs and with the rec. NS1 protein. Sera were then obtained from each group of animals and IgGs purified as described in the Materials and Methods. Naïve BALB/c mice (n=10 per group) were then transferred with 100 μg of the purified IgGs one day before and three days after an i.c. challenge with DENV2 NGC strain. Figure 5A shows the survival rates for each transferred group. No statistical difference was observed among the groups. Besides mortality, we also scored morbidity degrees (figure 5B) and, as observed in the survival rates, there was no statistical difference among the groups (p>0.05). These results suggest that, in our model, the anti-NS1 antibodies elicited by immunization with αDEC-NS1 or αDCIR2-NS1 mAbs do not mediate protection against and i.c. challenge.

**Figure 5.**
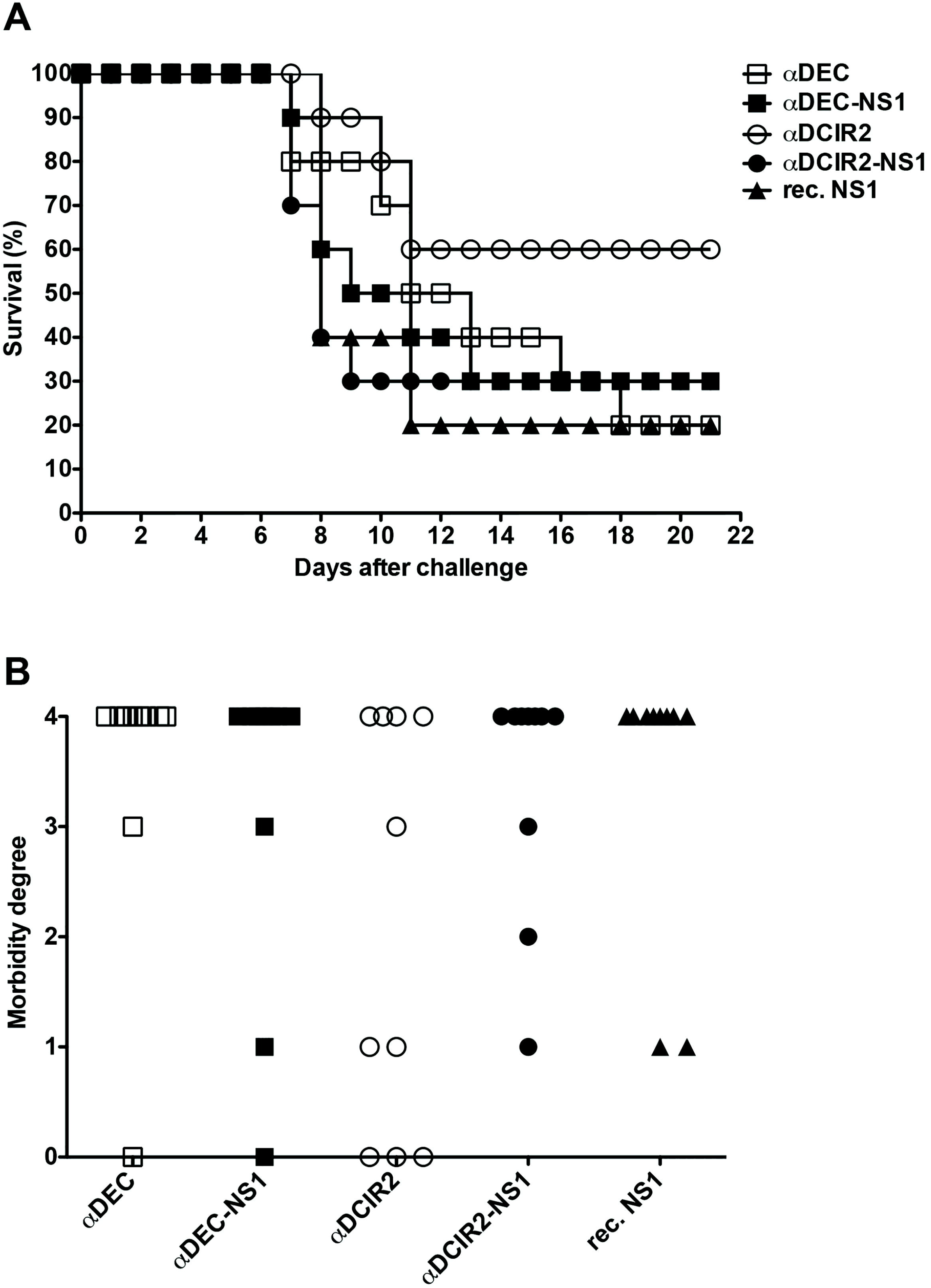

## Discussion

The controversy related to the induction of anti-NS1 antibodies and their binding to different cells and tissues prompted us to evaluate if these antibodies would be deleterious in our immunization context. For that purpose, we immunized groups of BALB/c mice with two hybrid mAbs fused to the NS1 protein (αDEC-NS1 or αDCIR2-NS1) and also one group with a recombinant NS1 protein produced in bacteria. As negative controls, groups were immunized with non-fused mAbs (αDEC or αDCIR2). We have previously shown that immunization with hybrid mAbs containing NS1 induced high anti-NS1 titres (17). To analyse these antibodies in more detail, we purified IgGs from the sera of immunized mice using protein G beads, and used the purified IgGs to perform binding assays to platelets. Only IgGs derived from animals immunized with αDCIR2-NS1 were able to bind very weakly to human platelets. None of the other purified IgGs presented any statistically significant binding when compared to the control groups. These results contrast with results obtained previously showing that anti-NS1 antibodies were able to bind to human platelets, more specifically to the protein disulfide isomerase (7, 8). This difference was surprising but may be explained by the fact that the authors hyper immunized a different mouse strain (C3H/HeN) with 5 immunizations using complete and incomplete Freunds adjuvant. In our case, BALB/c mice were immunized twice with only 5 μg of each hybrid mAb or rec. NS1 protein in the presence of poly (I:C). Despite the lower dose and the use of a different adjuvant, the anti-NS1 antibodies induced by immunization with both hybrid mAbs containing NS1 (αDEC-NS1 or αDCIR2-NS1) reacted very strongly with the rec. NS1 on ELISA plates, and a plateau was reached when we used the purified IgGs at concentrations ranging from 4 to 0.5 μg/mL. IgGs purified from mice injected twice with rec. NS1 showed a dose dependent binding curve, but proportionally ligated less well when compared to the purified IgGs obtained from mice immunized with the hybrid mAbs. When the avidity index was analysed, we were surprised by the fact that IgGs derived from rec. NS1 immunized mice presented the highest avidity, despite the fact they bound less to NS1. The biological meaning of such finding is difficult to predict.

The results obtained when we incubated different amounts of purified IgGs obtained from the different mouse groups directly with human platelets showed that only IgGs purified from the animals immunized with αDCIR2-NS1 were able to bind to a very small percentage of platelets and only when 5 μg were used. However, the analysis of platelet aggregation using these same purified IgGs revealed no interference in ADP-dependent aggregation, even with the antibodies generated by immunization with αDCIR2-NS1. Also, other parameters such as the bleeding time, platelet numbers or prothrombin time were not different among the groups. Thus, our results indicate that the antibodies against the NS1 raised in our immunization protocol do not seem to have any effect on platelet dysfunction. Again, these results contrast with those obtained by Chen et al. (7), and may be explained by the differences in mouse models and in immunization protocols.

Finally, we decided to transfer the purified IgGs transiently to naïve mice and challenge them subsequently. Once more, we did not observe any significant difference among the groups suggesting that the anti-NS1 antibodies induced by this immunization protocol do not protect or cause any other dysfunction that may accelerate death. In fact, we did not expect any protection mediated by anti-NS1 antibodies because we have previously shown that protection in this particular model is mediated by T cells (17). It is important to mention that protection mediated by CD4+ T cells and anti-NS1 antibodies was obtained when mice were immunized with a DNA plasmid encoding NS1. Interestingly, anti-NS1 antibodies were shown to participate in partial protection only when the intracranial challenge was performed using 4 LD50. If a higher dose was used (ie 40 LD50) protection was then mediated by the combination CD4+ T cells and anti-NS1 antibodies (15).

The relationship between the amount of circulating NS1 and dengue severity has already been established in humans (18, 23). The mechanism by which NS1 causes vascular leak was elucidated on a mouse model in two recent articles (4, 21). According to the authors, circulating NS1 is able to bind to the TLR4 receptor and induce: i) the release of proinflammatory cytokines and chemokines by PBMCs, and ii) vascular leakage in vitro and in vivo (21). Moreover, immunization with NS1 derived from DENV-1 to 4 was able to induce anti-NS1 antibodies that in turn protected mice against a lethal intravenous challenge with DENV2 (4). These previously published results, as well as the results described here, suggest that the anti-NS1 antibodies induced by different immunization protocols do not cause severe alterations that might be deleterious and induce pathology. However, in contrast to previously published data (4), the anti-NS1 antibodies induced by our immunization protocol did not protect BALB/c mice from an intracranial challenge with the DENV2 NGC strain. It is important to mention that the two models are quite different and protection obtained previously was reported against vascular leak and in *Ifnar*^*-/-*^ mice challenged with a different strain of DENV2.

Based on the results reported here, we conclude that targeting NS1 to two different DCs subsets induces high anti-NS1 antibody titres that do not bind significantly to platelets, block ADP-dependent platelet activation, cause any additional alteration in platelet numbers or increase bleeding tendency. Despite not deleterious, the anti-NS1 antibodies were also not able to protect mice against an intracranial challenge with DENV 2 NGC strain. The results described here provide evidence that anti-NS1 antibodies may have ambiguous roles and may not always lead to platelet dysfunction.

## Acknowledgements

This work was supported by the Brazilian National Research Council (CNPq, grant numbers 440721/2016-4), the Coordenação de Aperfeiçoamento de Pessoal de Nível Superior—Brasil (CAPES, Finance Code 001), the Decit/SCTIE/MoH and the São Paulo State Research Funding Agency (FAPESP, grant numbers 2013/11442-4 and 2014/50631-0). E.C.M.V received a post-doctoral fellowship from CAPES. S.B.B and D.S.R receive fellowships from CNPq.

We thank Dr. Beatriz Simonsen Stolf Carboni for critical reading of the manuscript; Danielle Chagas, Juliane Pereira Afonso, Anderson Antonio Domingos Silva and Jonatas Pereira for assistance at the animal facility.

